# Flexibility of territorial aggression in urban and rural Chaffinches

**DOI:** 10.1101/2025.10.29.685300

**Authors:** Alper Yelimlieş, Çağla Önsal, Çağlar Akçay

## Abstract

Rapid environmental change due to urbanization poses novel challenges to animals. Behavioral change and individual plasticity are generally hypothesized to be the key to adapting to these challenges. One commonly observed behavioral change is higher observed aggression levels in urban animals, perhaps because anthropogenic noise disrupts effective acoustic communication during conflicts, leading to greater use of physical aggression. We investigated the hypothesis that urban noise drives aggression by performing repeated simulated territorial intrusion experiments on rural and urban chaffinches (*Fringilla coelebs*). We expected urban chaffinches to be more aggressive, change their aggression levels more between trials, and for aggression to increase with noise levels, irrespective of the habitat. We found that while aggression didn’t differ between habitats in the initial trial, rural chaffinches decreased their aggression level in the second trial and thus were less aggressive than the urban chaffinches, who did not change their response. That is, urban birds were less flexible in responding to an intruder than rural birds, contrary to previous findings in other songbirds. Moreover, there was an increase in aggression with increasing noise levels, irrespective of the habitat. Given our small sample size and lack of spatial replicates, our results should be interpreted with caution. Nevertheless, as a lack of flexibility in aggression is potentially costly, our results highlight the importance of studying the plasticity in aggressive behavior in human-impacted landscapes.

---

Urbanization presents novel and varied evolutionary challenges to animals, including pollution, habitat fragmentation, and climate change (Grimm et al., 2008). Populations facing these challenges in cities undergo rapid evolution as a result of adaptive and nonadaptive processes (Johnson & Munshi-South, 2017; Sih et al., 2011). These urban dwellers often shift their behavioral phenotypes to survive in the human-impacted environments, making behavioral change a precursor to urban evolution (Caspi et al., 2022; Sol et al., 2013). Indeed, differences in behavioral traits between rural and urban populations of animals are well documented (Gil & Brumm, 2014; Lowry et al., 2013; Ritzel & Gallo, 2020). Compared to their rural counterparts, urban-dwelling individuals of a species are shown to be bolder (Samia et al., 2015) and less aversive to novelty (Griffin et al., 2017; Tryjanowski et al., 2016), to breed earlier (Capilla□Lasheras et al., 2022), and to differ in their signaling behaviors (Slabbekoorn & Peet, 2003). One important phenotypic change associated with urbanization is that urban animals tend to be more aggressive compared to rural animals (Abolins□Abols et al., 2016; Colombelli-Négrel et al., 2023; Davies & Sewall, 2016; Diniz & Duca, 2021; Fokidis et al., 2011; Foltz et al., 2015; Hardman & Dalesman, 2018; Önsal et al., 2022; but see Hasegawa et al., 2014; Hurtado & Mabry, 2017).

It is unclear why urban animals tend to be more aggressive than their rural counterparts, although several explanations have been proposed. These include increased food availability (Foltz et al., 2015), behavioral syndromes (Evans et al., 2010), metal pollution (McClelland et al., 2019), and noise pollution (Phillips & Derryberry, 2018). Among these factors, the latter has attracted significant research attention (Akçay et al., 2020a; Diniz & Duca, 2021; Grabarczyk & Gill, 2019; Önsal et al., 2022). It is hypothesized that anthropogenic noise contributes to increased aggression because signals become less efficient due to masking, which prevents the resolution of disputes through acoustic communication, ultimately leading to physical fights (Akçay et al., 2020b; De Kort et al., 2024).

Behavioral flexibility can allow individuals to cope with rapid environmental changes and so is hypothesized to be a key component of urban adaptation (Caspi et al., 2022; Sih et al., 2011). Experimental studies showed that individuals dynamically increase their physical aggressive behavior when presented with noise (De Kort et al., 2024; Grabarczyk & Gill, 2019; Hohl et al., 2025; Önsal et al., 2022). Likewise, dark-eyed juncos (*Junco hyemalis*) living in urban habitats showed decreased aggression towards conspecifics during the COVID-19 pandemic (Walters et al., 2022), presumably as a result of the concurrent decrease in human activities during successive lockdowns (“anthropause”; Rutz et al. 2020). Again, these results suggest individual flexibility in response to noise in urban habitats. Consequently, as within-individual variation increases, repeatable individual differences may decrease in urban habitats. A study with great tits (*Parus major*) supported this idea. Hardman and Dalesman (2018) found significant repeatability of all five proxies of aggression for rural birds, but for only two of them for urban birds, although their results were inconclusive, as there were no differences in repeatability estimates for each variable between urban and rural populations.

Here, we studied differences in the intensity and repeatability of territorial aggression between urban and rural populations of common chaffinch (*Fringilla coelebs*), particularly evaluating the anthropogenic noise hypothesis. Using song playback, we simulated conspecific intrusions inside male territories on two consecutive days. First, we predicted that territorial aggression would be higher in urban chaffinches, consistent with previous studies comparing urban and rural animal populations. Because we hypothesized that anthropogenic noise can disrupt acoustic communication, we also predicted that ambient noise levels would correlate positively with aggression irrespective of habitat type. Moreover, we reasoned that mostly aggressive individuals, and those that can flexibly adjust to fluctuating environmental conditions, would be better suited to urban habitats. Hence, we predicted that urban chaffinches would have lower among-individual variance, higher within-individual variance, and lower repeatability with respect to their aggressiveness.

## Methods

### Study site and species

We chose chaffinches for investigating differences with regard to territorial aggression, as their territorial behaviors are well studied and they are frequently found in both urban and rural habitats (Brumm & Ritschard, 2011; Marler, 1956; Slater, 1981). Chaffinches are territorial only during the breeding season; territories are established by males, which are about 0.7 ha (Marler, 1956). Males defend their territories from intrusions. In the case of playback experiments, defense typically involves flights around the speaker but not singing, which suggests song is used as a keep-out signal but not in active defense (Slater, 1981).

We performed simulated territory intrusion experiments on 23 male chaffinches inhabiting one urban site (n = 11) and two rural sites (n = 12) in Sarıyer, İstanbul, Turkey. Ambient noise in the urban site was significantly higher than in the rural sites, making the locations suitable for our investigation (Yelimlieş et al., 2023). In our study, each male was tested twice; however, none of the subjects were ringed in the study, so we assumed the territory holders did not change between the two consecutive trials, which were separated by 24 hours. We avoided including neighboring males in our sample and did not test males located closer than 200 meters to each other in order to ensure that each male is included in our sample only once. All trials were conducted between 11 and 26 May 2021.

### Playback Stimuli

For the simulated territory intrusions, we used songs of chaffinches between March and May 2019 along with songs of the subjects in this study, which were recorded using a Marantz PMD660 or 661 recorder with a Sennheiser ME66/K6 shotgun microphone. From our recordings, we selected 22 high quality songs from 22 different male chaffinches. We removed low-frequency noise from each song recording using a high-pass filter (threshold 1000 Hz) in Raven Pro 1.6.1 (K. Lisa Yang Center for Conservation Bioacoustics, 2019). Then we added silence at the end of each song to create 10-second-long stimulus files. The mean ± SD song duration was 2.27 ± 0.27 secs.

### Experimental Procedure

We found and observed territorial chaffinches in our sites to determine the singing posts of the focal and neighboring individuals. Prior to song playback, we placed a wireless speaker (Anker SoundCore, Anker, Inc.) about 2 m above the ground level, and within the boundaries of the singing posts of the resident male for playing the conspecific stimulus. After positioning ourselves away from the speaker and confirming the presence of the resident male within the territory, we played the conspecific song stimulus continuously 6 times (1 minute playback in total). During this playback period, we narrated the flights of the focal male, noting the distance from the speaker. We used either a Marantz PMD660 recorder with a Sennheiser ME66/K6 shotgun microphone or a Zoom H5 recorder with a Zoom SGH-6 shotgun microphone to record our narrations. After each trial, we measured the average ambient noise level in dB in the target male’s territory using a sound level meter (VLIKE VL6708, VLIKE Inc., A-weighted, fast). To get an average, we obtained 8 noise levels measuring twice from 4 directions that are perpendicular to each other from a single location in the territory (Brumm, 2004).

### Data Analysis

Using Raven Pro 1.6.1, we annotated our narrations of the trial and coded the proportion of time spent within 1 and 5 meters of the speaker, number of flights, and closest approach to the speaker for each trial. These variables are used as proxies of aggression levels in songbirds (Brumm & Ritschard, 2011), and have been shown to predict attacking on a taxidermic model (Akçay et al., 2013). We then performed a principal component analysis (PCA) to obtain a composite score for aggression. We evaluated the significance of PCA and principal components using the *PCAtest* package with 1000 permutations and bootstrap replication (Camargo, 2022). Permutation tests showed that only PC1 was meaningful, which accounted for 71.5% (95%-CI = 61.6-79.5) of the total variation in the data, and had an eigenvalue of 2.86. All variables had significant loadings on PC1 (number of flights = 0.47, closest approach = -0.55, time spent within 5m = 0.47, and 1m = 0.50). So, we ran the PCA using the function *prcomp* from the *stats* package with centered and scaled variables and took PC1 scores as the aggression scores.

We then modeled the relationship between aggression scores and habitat type via a linear mixed model with Gaussian error distribution using the function *lmer* from the package *lme4* (Bates et al., 2015). As exploratory visualizations showed a discrepancy between first and second trials depending on habitat, the model included habitat, trial order, and their interaction effect as fixed factors and male ID as the random factor. We estimated the repeatability for ambient noise and aggressiveness, as well as between- and within-individual variance for aggressiveness, separately for each habitat, using the package *rptR* (Stoffel et al., 2017), with a Gaussian error distribution and calculated 84% confidence intervals using bootstrapping with 1000 iterations. We chose 84% because non-overlapping 84% confidence intervals can be used as a proxy of the difference in estimates being different than zero at α = 0.05 (Payton et al., 2003).

Lastly, we modeled the relationship between aggression scores and ambient noise levels while controlling for habitat, trial order, and their interaction, using another linear mixed model with a Gaussian error distribution. The model included male ID as a random effect. A subset of males (4 rural and 1 urban) had missing ambient noise measurements for one trial from their territory and thus were excluded from this analysis, reducing the sample size for to 41 trials from 23 males. Given the almost complete separation of noise levels between habitats (see Results), we then repeated this analysis with the urban subset (N = 11 males, 18 trials) with trial order as the fixed effect and male ID as the random effect. We reasoned that, since noise levels vary more in the urban habitat, if there is a relationship between noise and aggression, this would strengthen our conclusion that the relationship is independent of the habitat.

We checked multicollinearity for both models by calculating variance inflation factors using the function *vif* from the package *car* (Fox & Weisberg, 2019), confirming that all values were < 2.4. We checked residual diagnostics using the package DHARMa (Hartig, 2024). We report coefficient estimates, standard errors from the output of *lmer* function, as well as χ*2* and *P* values for evaluating statistical significance for the overall terms using the *Anova* function from the package *car*. To interpret the interaction effect, we performed post-hoc tests with Tukey-adjusted *P* values using the *pairs* function from the package *emmeans* (Lenth, 2024). We plotted our data using the packages *ggplot2* (Wickham, 2016) and *ggeffects* (Lüdecke, 2018).

All the statistical analyses are performed in R (R Core Team, 2024). Details of package versions and dependencies can be found at the end of the R script provided in the supplementary materials.

## Results

Aggressive behavior during simulated territory intrusions depended on the interaction between habitat and trial order (interaction effect: χ*2* = 6.24, *P* = 0.01, see Table 1, Figure 1). In their first trials, aggression scores did not differ between urban and rural chaffinches (contrast estimate = -0.97, SE = 0.61, *P* = 0.39). However, chaffinches in rural habitats became less aggressive in their second trial (contrast estimate = 1.22, SE = 0.35, *P* = 0.01) while urban chaffinches showed no change in aggression between trials (contrast estimate = -0.06, SE = 0.37, *P* = 0.99). As a result, urban chaffinches had higher aggression scores in their second trial than rural chaffinches (contrast estimate = -2.25, SE = 0.61, *P* = 0.004).

**Table 1.** Output from a linear mixed model assessing whether aggression scores are influenced by habitat (urban, rural) and trial order (first, second). Bold values indicate statistical significance (P < 0.05).

| *Predictor* | *Estimate* | *95 % CI* | $\chi^2$ | *df* | *P* |
| --- | --- | --- | --- | --- | --- |
| Intercept | -0.16 | -1.02 – 0.69 | 0.15 | 1 | 0.70 |
| habitat [urban] | 0.97 | -0.26 – 2.21 | 2.54 | 1 | 0.11 |
| **trial order [second]** | **-1.22** | **-1.93 – -0.50** | **11.81** | **1** | **<0.001** |
| **habitat [urban] x trial order [second]** | **1.28** | **0.24 – 2.31** | **6.24** | **1** | **0.01** |

**Figure 1.**
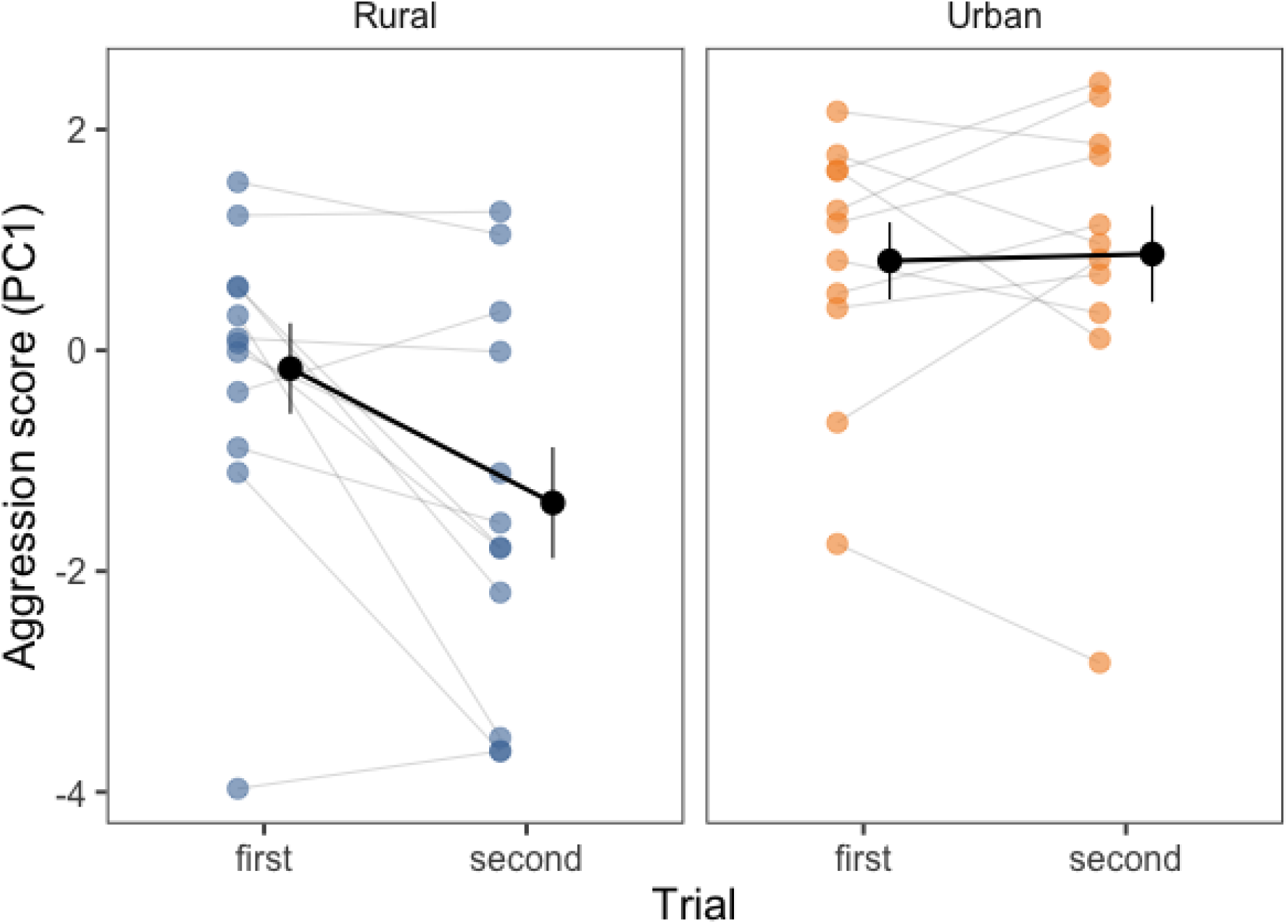
Aggression scores (PC1, higher values indicate higher aggression) in response to a simulated territory intrusion in both urban and rural chaffinches. Although rural chaffinches decreased their aggressive response on their second encounter with a simulated intruder, urban chaffinches did not. Each coloured circle represents a single trial, with individuals’ first and second trials connected by a line. Group means and standard errors are represented with the darker circles and vertical lines to their right.

Aggression across the two playback trials was highly repeatable in urban chaffinches (R = 0.75, 84% CI = 0.47–0.89, *P* = 0.006) but was not significantly repeatable in rural chaffinches, although the estimates didn’t differ from each other (R = 0.41, 84% CI = 0.02–0.70, *P* = 0.087). Urban chaffinches had significantly lower within-individual variance than rural chaffinches (urban = 0.42, 84% CI = 0.19–0.67; rural = 1.68, 84% CI = 0.81–2.57), while among-individual differences did not differ between habitats (urban = 1.28, 84% CI = 0.49–2.37; rural = 1.18, 84% CI = 0–2.5).

Ambient noise levels were significantly repeatable in both habitats (Urban: R = 0.78, 84% CI = 0.56-0.91, *P* = 0.001; Rural: R = 0.71, 84% CI = 0.36-0.89, *P* = 0.016). Moreover, controlling for habitat, trial order, and their interaction, the effect of ambient noise levels on aggression scores was significant (estimate = 0.11, SE = 0.05, χ*2* = 4.00, *P* = 0.046). Yet, a separate analysis with the urban subset revealed that there was no relationship between noise and aggression (estimate = 0.09, SE = 0.33, χ*2* = 3.13, *P* = 0.08), where noise levels were more variable (see Figure 2).

**Figure 2.**
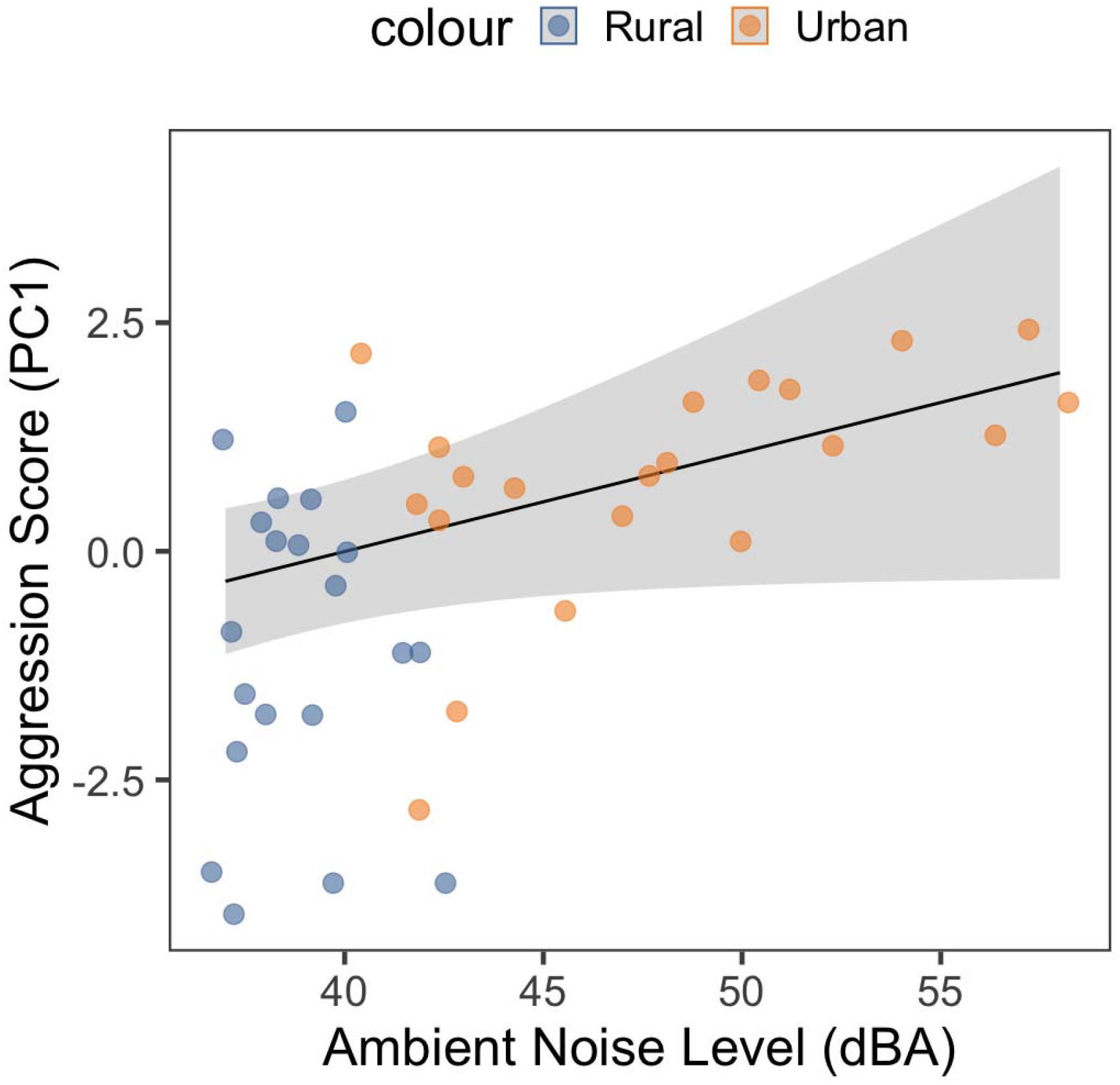
Scatterplot showing the positive correlation between aggression score (PC1) in male chaffinches and the ambient noise level (dBA) in their rural or urban territory. There was no such relationship when the urban subset was inspected alone. The points show each trial, and the line shows predicted values controlling for other fixed and random effects in the model, and the gray shading shows 95% CI.

## Discussion

In this study, we tested whether the intensity and flexibility of aggressiveness differed between urban and rural chaffinches in response to simulated territory intrusions, and also whether anthropogenic noise predicts aggression irrespective of habitat. We found that habitat differences in aggression depended on trial order. Although urban and rural chaffinches did not differ in the first trial, urban birds were more aggressive than their rural counterparts in the second trial. Rural birds reduced their aggressiveness in their second trial while urban birds did not. Consequently, urban chaffinches were less flexible in aggression. Although we found a significant positive correlation between ambient noise and aggression scores, this relationship was absent in the urban site alone. Therefore, our data are not appropriate for making strong conclusions about the effect of noise on aggression due to the almost non-overlapping distribution of noise between the habitats.

As we predicted, urban chaffinches were more aggressive than rural chaffinches, albeit only during their second territory intrusion trial. Many prior studies have reported higher overall aggressiveness among urban birds (Davies & Sewall, 2016; Hardman & Dalesman, 2018; Önsal et al., 2022; but see Bókony et al., 2010). However, Beck et al. (2023) reported a similar interaction between habitat and trial number, whereby rural and urban song song sparrows (*Melospiza melodia*) did not differ in aggressiveness during their first simulated territorial intrusion, but urban birds became more aggressive than rural birds in their next trial two weeks later.

In our study, urban chaffinches tested on consecutive days did not change their aggression levels across trials, while rural chaffinches decreased their aggression on the second day. This may be indicative of several different processes. Our design, which used the same stimuli across two trials for each male, may have enhanced this effect. While this design avoids confounds in which specific stimulus characteristics may drive responses, it also may have been perceived by the subjects as a previously repelled male intruding again. Rural chaffinches may have reduced their territorial effort based on that previously-won contest, while urban birds maintained their original effort level. This might have been caused by interrupted individual recognition in the urban habitat due to noise. An alternative hypothesis is that reducing aggression might be too costly for urban chaffinches if territory resource value is higher in urban habitats, due to more clumped habitat and resource distribution (Foltz et al., 2015; Marzluff 2001; Juárez et al. 2020).

Whatever the cause, it seems urban birds show reduced within-individual variation in their behavior compared to rural birds. This finding is inconsistent with the hypothesis that urban adapters should have greater behavioral flexibility (Caspi et al., 2022). Nevertheless, other studies investigating different bird species reported similar findings (e.g. Beck et al, 2023). In dark-eyed juncos, stress causes a decrease in the aggressiveness of rural but not urban dark-eyed juncos (Abolins□Abols et al., 2016). Conversely, rural but not urban European robins (*Erithacus rubecula*) increased aggression when presented with experimental noise (Önsal et al., 2022). As being able to adjust aggression would prevent unnecessary injuries or energy wastage, behavioral inflexibility may have serious fitness costs for urban animals. Although our results highlight the importance of studying the flexibility of aggressive behavior in human-impacted environments, these results should be interpreted with caution because our sample size was limited and we were only able to study chaffinches in one urban habitat.

## Acknowledgements

We thank Andrew C. Katsis for his thorough feedback on the manuscript. This project is partly funded by the Austrian Science Fund (Project numbers 10.55776/W1262 and 10.55776/P36342) with awards to Sonia Kleindorfer.

## Data availability

Data and R script to reproduce analyses can be found in the supplementary materials.

## Author contributions

AY and CA designed the research; AY, CO, and CA conducted the playback trials. AY analyzed the data and wrote the first draft of the manuscript; all authors revised the manuscript and gave the final approval for publication.

